# Revealing 29 sets of independently modulated genes in *Staphylococcus aureus*, their regulators and role in key physiological responses

**DOI:** 10.1101/2020.03.18.997296

**Authors:** Saugat Poudel, Hannah Tsunemoto, Yara Seif, Anand Sastry, Richard Szubin, Sibei Xu, Henrique Machado, Connor Olson, Amitesh Anand, Joe Pogliano, Victor Nizet, Bernhard O. Palsson

## Abstract

The ability of *Staphylococcus aureus* to infect many different tissue sites is enabled, in part, by its Transcriptional Regulatory Network (TRN) that coordinates its gene expression to respond to different environments. We elucidated the organization and activity of this TRN by applying Independent Component Analysis (ICA) to a compendium of 108 RNAseq expression profiles from two *S. aureus* clinical strains (TCH1516 and LAC). ICA decomposed the *S. aureus* transcriptome into 29 independently modulated sets of genes (i-modulons) that revealed (1) high confidence associations between 21 i-modulons and known regulators; (2) an association between an i-modulon and σS, whose regulatory role was previously undefined; (3) the regulatory organization of 65 virulence factors in the form of three i-modulons associated with AgrR, SaeR and Vim-3, (4) the roles of three key transcription factors (codY, Fur and ccpA) in coordinating the metabolic and regulatory networks; and (5) a low dimensional representation, involving the function of few transcription factors, of changes in gene expression between two laboratory media (RPMI, CAMHB) and two physiological media (blood and serum). This representation of the TRN covers 842 genes representing 76% of the variance in gene expression that provides a quantitative reconstruction of transcriptional modules in *S. aureus*, and a platform enabling its full elucidation.

**Significance Statement:** *Staphylococcus aureus* infections impose an immense burden on the healthcare system. To establish a successful infection in a hostile host environment, *S. aureus* must coordinate its gene expression to respond to a wide array of challenges. This balancing act is largely orchestrated by the Transcriptional Regulatory Network (TRN). Here, we present a model of 29 independently modulated sets of genes that form the basis for a segment of the TRN in clinical USA300 strains of *S. aureus*. Using this model, we demonstrate the concerted role of various cellular systems (e.g. metabolism, virulence and stress response) underlying key physiological responses, including response during blood infection.

## Introduction

*Staphylococcus aureus* causes a variety of human diseases ranging from skin and soft tissue infections (SSTI) to infective endocarditis and pneumonia^1^. The pathogen can also thrive as part of the commensal microbiome in the anterior nares of healthy patients^2^. *S. aureus* adaptation to many different host environments is enabled, in part, by the underlying transcriptional regulatory network (TRN) that can alter the physiological state of the cell to match the unique challenges presented by each environment^3–5^. Such adaptations require coordinated expression of genes in many cellular subsystems such as metabolism, cell wall biosynthesis, stress response, virulence factors, etc. Therefore, a complete understanding of the *S. aureus* response to different environments necessitates a thorough understanding of its TRN. However, since *S. aureus* is predicted to have as many as 135 transcriptional regulators^6^, with many more potential interactions among them, a bottom-up study of its global TRN becomes intractable.

To address this challenge, we previously introduced an Independent Component Analysis (ICA)-based framework in *Escherichia coli* that decomposes a compendium of RNA-sequencing (RNA-seq) expression profiles to determine the underlying regulatory structure^7^. An extensive analysis of module detection methods demonstrated that ICA out-performed most other methods in consistently recovering known biological modules^8^. The framework defines independently modulated sets of genes (called i-modulons) and calculates the activity level of each i-modulon in the input expression profile. ICA analysis of expression profiles in *E. coli* have been used to describe undefined regulons, link strain-specific mutations with changes in gene expression, and understand rewiring of TRN during Adaptive Laboratory Evolution (ALE)^7,9^. Given the deeper insights it provided into the TRN of *E. coli*, we sought to expand this approach to the human pathogen *S. aureus*.

To elucidate the TRN features in *S. aureus*, we compiled 108 high quality RNA-seq expression profiles for community-associated methicillin-resistant *S. aureus* (CA-MRSA) strains LAC and TCH1516. Decomposition of these expression profiles revealed 29 independently modulated sets of genes and their activity levels across all 108 expression profiles. Further, we show that using the new framework to reevaluate the RNA-seq data accelerates discovery by (1) quantitatively formulating TRN organization, (2) simplifying complex changes across hundreds of genes into a few changes in regulator activities, (3) allowing for analysis of interactions among different regulators, (4) connecting transcriptional regulation to metabolism, and (5) defining previously unknown regulons.

## Main

### ICA extracts biologically meaningful components from transcriptomic data

We generated 108 high-quality RNA-seq expression profiles from CA-MRSA USA300 isolates LAC and TCH1516 and two additional Adaptive Laboratory Evolution (ALE)-derivatives of TCH1516. To capture a wide range of expression states, we collected RNA-seq data from *S. aureus* exposed to various media conditions, antibiotics, nutrient sources, and other stressors (see **Supplementary Table 1**). The samples were then filtered for high reproducibility between replicates to minimize noise in the data (Figure S1a). The final dataset contained 108 samples representing 43 unique growth conditions, which have an average R^2^ = 0.98 between replicates.

Using an extended ICA algorithm^7^, we decomposed the expression compendium into 29 ‘i-modulons’. An i-modulon contains a set of genes whose expression levels vary concurrently with each other, but independently of all other genes not in the given i-modulon. Akin to a regulon^10^, an i-modulon represents a regulatory organizational unit containing a functionally related and co-expressed set of genes under all conditions considered (**Figure 1a**). While regulons are determined based on direct molecular methods (e.g., ChIP-seq, RIP-ChIP, gene-knockouts, etc.), i-modulons are defined through an untargeted ICA-based statistical approach applied to RNA-seq data that is a reflection of the activity of the transcriptional regulators (see Materials and Methods). However, beyond regulons, i-modulons can also describe other genomic features, such as strain differences and genetic alterations (e.g. gene knock-out) that can lead to change in gene co-expression^9,11^. The outcome of this approach is a biologically relevant, low dimensional mathematical representation of functional modules in the TRN that reconstruct most of the information content of the input RNAseq compendium (**Figure S1b**). Such formulation also quantitatively captured complex behaviors of regulators such as contra-regulation of multiple genes by the same regulator, co-regulation of the same gene by multiple regulators, and coordinated expression of multiple organizational units (i-modulons) in various conditions (**Figure S1c-d**). Therefore, this model enables simultaneous analysis of TRN at both gene and genome-scale.

**Figure 1:**
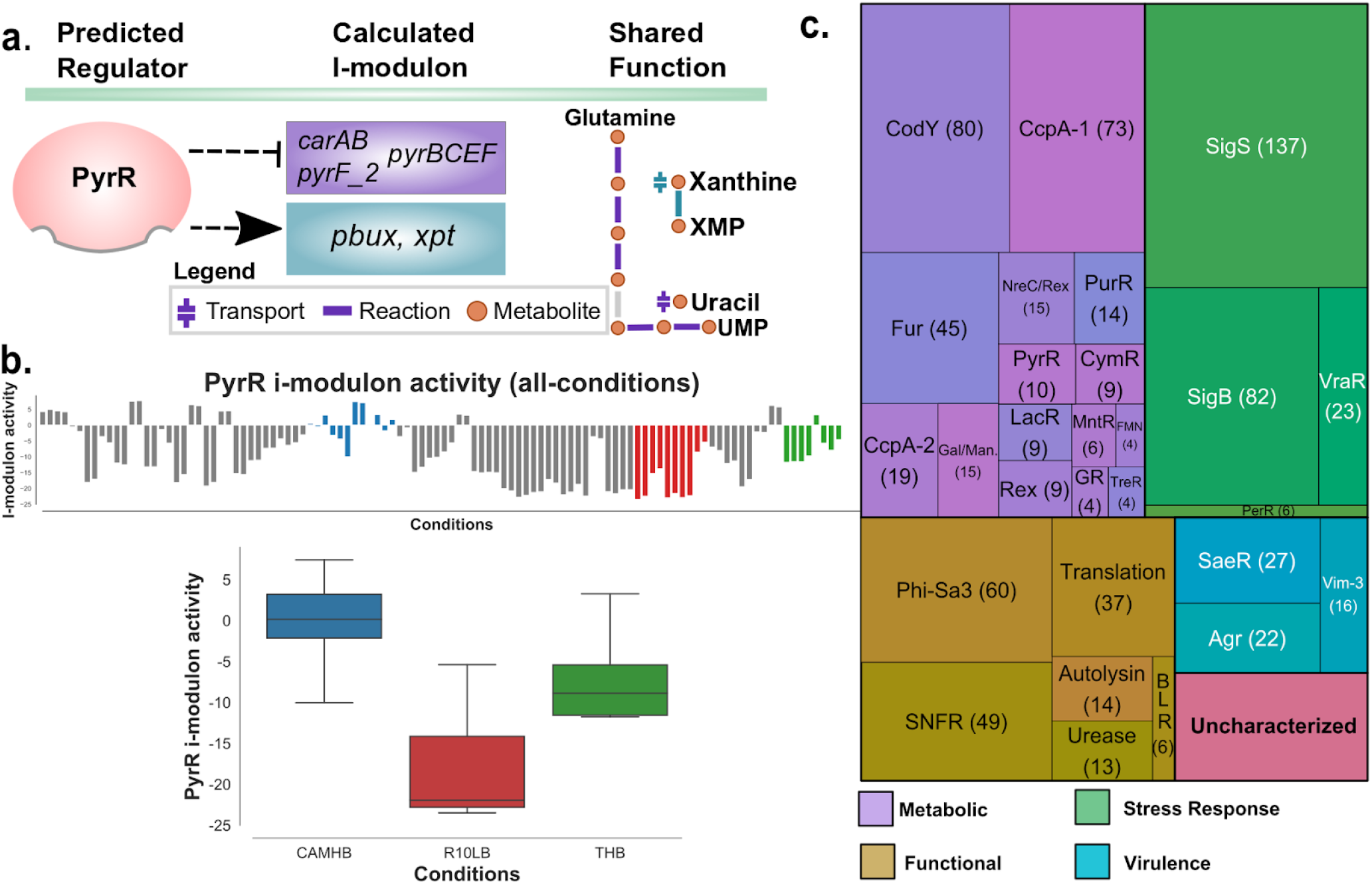
ICA decomposition of *S. aureus* USA300 RNA-seq database. (a) An i-modulon is a set of genes that are co-expressed and encode products with shared functions. The PyrR i-modulon, for example (middle column), is predicted to be under control of pyrR repressor and contains genes that encode enzymes in pyrimidine biosynthesis (purple) and purine salvage (blue) pathway (right column). The genes in two different pathways are contra-regulated (arrows). (b) Activity levels of i-modulons are calculated for all conditions (top bar chart), allowing for sample specific (e.g., in three different media) comparison of each i-modulon (boxplot). The activity of all i-modulons are centered around CAMHB base condition and therefore, all i-modulons have mean activity of 0 in this condition. Centerline of the boxplot represents median value, the box limits represent Q1 and Q3, and the whiskers represent the min and max values. (c) A treemap indicating the names and the size of the i-modulons. The i-modulons are named after the transcription factor(s) whose predicted regulons have highest overlap with the given i-modulon, or based on the shared functionality of genes (e.g., autolysins, translation, B-lactam resistance) in i-modulon if no known regulator was identified. i-modulon with low or no correspondence with any of the known features is labeled as Unc-1. ‘BLR’ stands for ‘Beta-Lactam resistance’ and SNFR i-modulon consists of genes with altered expression in SNFR strain.

ICA also reconstructs the activity of the i-modulons in the samples, which represents the collective expression level of the genes in the i-modulon. Each sample in the dataset can be reconstructed as the summation of the activity of the 29 i-modulons, which makes the transcriptional state in each condition more explainable. Conversely, each i-modulon has a computed activity in every sample, allowing for easy comparisons of i-modulon activities across samples, that in turn reflect the activity of the corresponding transcriptional regulator (**Figure 1b**). The reported activity levels are log2 fold change from the base condition - growth in Cation Adjusted Mueller Hinton Broth (CAMHB).

We compared the gene sets in the 29 enriched i-modulons against previously predicted \textit{S. aureus} regulons in the RegPrecise database and other regulons described in various publications (Supplementary Table 2). I-modulons with statistically significant overlap (FDR < 1e-05) with a previously predicted regulon were named after the transcription factor associated with the regulon (see Materials and Methods). We also manually identified i-modulons that consisted of genes with shared functions (e.g., Autolysins, Translation) or those that corresponded to other genomic features such as plasmids, prophages, or strain-specific differences. Together, we identified fifteen metabolic, six functional, three virulence, four stress response-associated and one strain-associated (SNFR) i-modulons (Figure 1c, Supplementary Table 3). Of the 29 enriched i-modulons, only one remains uncharacterized (See Supplementary Table 4). In total, the 29 i-modulons consist of 752 unique genes, 90 of which are enriched in more than 1 i-modulons.

### ICA disentangles complex change in the transcriptome

Differential expression analysis of *S. aureus* in different environmental conditions can yield hundreds of genes that have significantly altered expression levels, hindering meaningful interpretation. Decomposition of the expression profile into biologically meaningful i-modulons instead allows us to gain a comprehensive understanding of the change in the transcriptome through the activities of few regulators. To demonstrate this capability, we explored the difference in expression profiles of *S. aureus* grown in two different media, cation-adjusted Mueller Hinton Broth (CAMHB), the standard bacteriologic medium for routine antimicrobial susceptibility testing worldwide, and the common physiologically relevant mammalian tissue culture medium RPMI-1640, supplemented with 10% Luria broth (R10LB) to support growth kinetics similar to CAMHB.

Over 800 genes spanning more than a dozen Clusters of Orthologous Groups (COG)^12^ categories were differentially expressed between the two media (**Figure S2a**). Conversely, there were fifteen i-modulons with statistically significant differential activation **(Figure 2a**). Most differentially activated i-modulons were involved in metabolism (CodY, PurR, Guanine-Responsive i-modulon (GR), Gal/Man, Rex, MntR, PyrR, LacR, CcpA-1, CcpA-2, Urease). The last four i-modulons were those with functions in virulence (Vim-3, SaeR), translation, and the Phi-Sa3 phage-specific i-modulon.

**Figure 2:**
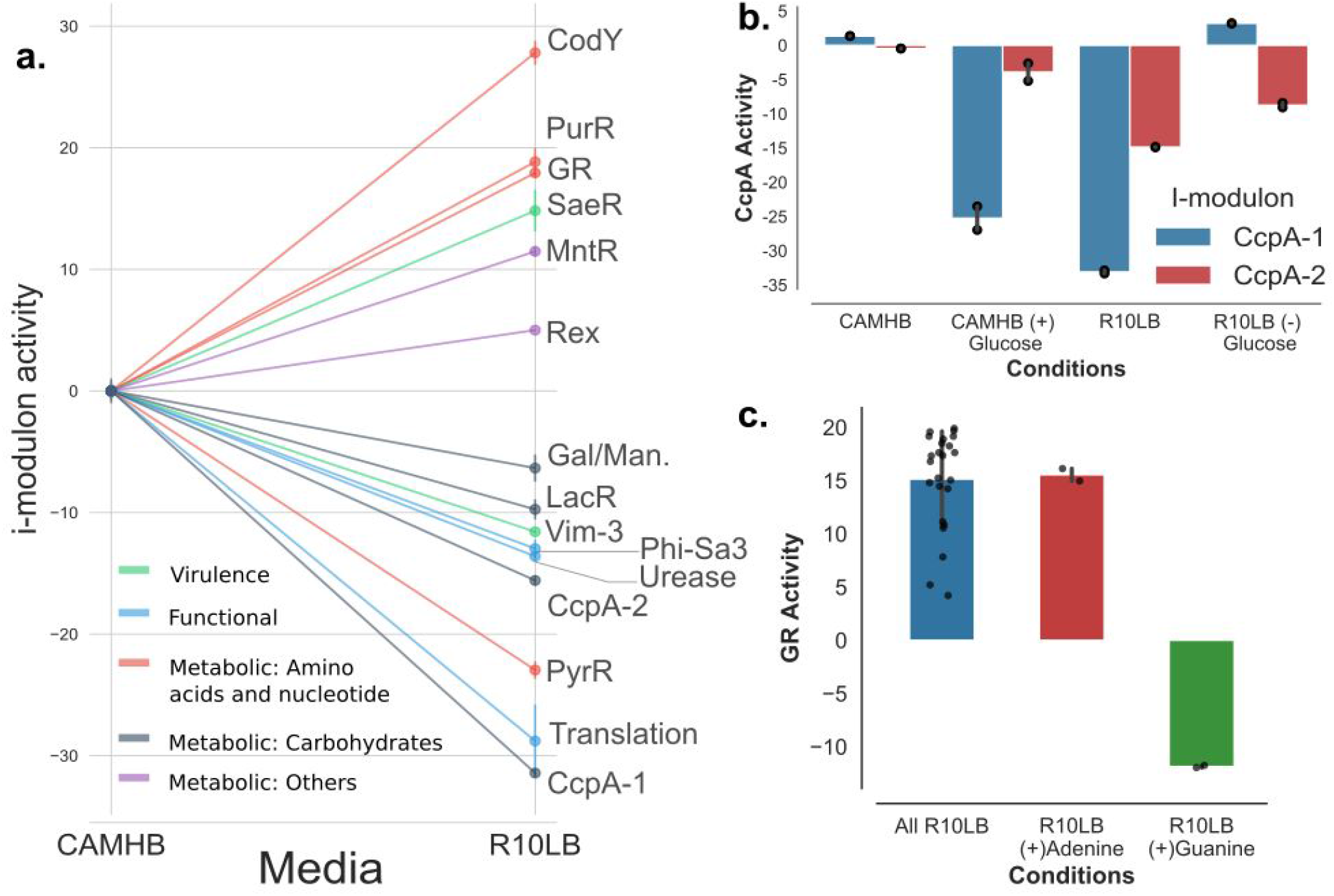
Differential activation of i-modulon in different media conditions. (a) I-modulons from LAC strain with statistically significant (p-value < 0.05) differential activation in CAMHB versus R10LB. (b) Addition of glucose reduced the activity of CcpA-1 in CAMHB (blue bars). Conversely, replacing glucose with maltose led to higher CcpA-1 activity in R10LB. CcpA-2 activity did not change in response to glucose concentration (red bars). (c) The bar plot shows the activity level of GR i-modulon, which contains the genes under the control of guanine riboswitch (*xpt and pbuG*). Though many different conditions can affect the GR i-modulon activity (blue bar), it sharply decreases when guanine is added to the media. The change in GR activity is guanine specific, as addition of adenine has no effect. Black dots in panel b and c represent values from individual samples and error bars represent standard deviation.

Concurrent activation of the CodY, PurR, and GR i-modulons in R10LB indicates that this media presents a guanine-limited environment, as activity of all three transcription factors decrease in response to falling cellular concentrations of various forms of guanine derivatives^13–16^. Consistent with this hypothesis, we also saw decreased activity of the Translation i-modulon in R10LB. Downregulation of translation machinery often occurs during the stringent response, where cellular GTP is depleted as it is rapidly converted to ppGpp^13,17–19^. Similarly, activation of the MntR i-modulon points to manganese starvation in R10LB, and the decreased activity of two i-modulons associated with carbon catabolite repressor CcpA (CcpA-1 and CcpA-2) likely reflects a glucose replete environment^20^. Analysis of spent media using HPLC confirmed that *S. aureus* was actively uptaking glucose in R10LB while no glucose was detected in CAMHB (**Figure S2b**). Taken together, the shift in activity of i-modulons between the two media suggests that compared to the bacteriologic medium CAMHB, R10LB presents an environment poor in purines (specifically guanine) and manganese but rich in the carbon source glucose.

Next, we designed two validation experiments to ensure that the activity level of i-modulons reflect expected outcomes. To this end, we chose three i-modulons to validate, CcpA-1, CcpA-2, and GR, for ease of modifying their activities with supplementation of glucose and purines, respectively. CcpA is the carbon catabolite repressor in *S. aureus* that controls central carbon metabolism and carbon source utilization^21,22^. Its activity level is indirectly modulated by cellular glucose concentration, though it can also be altered by other glucose-independent signals^23,24^. CcpA transcriptional effects are captured in two i-modulons, CcpA-1 and CcpA-2, which contain 73 and 19 genes, respectively. Both i-modulons had far lower activity in R10LB compared to CAMHB. However, the addition of 2g/L glucose only led to reduced activity of the CcpA-1 i-modulon in CAMHB, closely matching its activity in R10LB (**Figure 2b**). Similarly, replacement of glucose with maltose in R10LB led to increased activity of the CcpA-1 i-modulon. The change in glucose concentration, however, had little effect on the activity level of the CcpA-2 i-modulon, suggesting that the CcpA-1 i-modulon represents direct glucose-responsive CcpA activity, whereas the CcpA-2 i-modulon may reflect its glucose-independent activity.

In addition to CcpA activity, we also confirmed the activity of the GR i-modulon. The GR i-modulon contains genes involved in the purine salvage pathway (*xpT, pbuX)*, peptide transport (*oppB)*, and LAC specific virulence factor *ssl11*. The two genes in the salvage pathway have been previously demonstrated to be under the control of the guanine riboswitch in *S. aureus* strain NRS384^16^. The presence of this riboswitch was confirmed using the online RiboSwitch Finder (**Figure S2c-d**)^25^; no such riboswitch was detected for the other two genes. The activity of the i-modulon was attenuated by guanine supplementation (25ug/mL) while the addition of adenine had no effect, demonstrating a guanine-specific activity of the i-modulon (**Figure 2c**).

### Integration of i-modulons with genome-scale models of metabolism reveal systems-level properties of metabolic regulation

Genome-scale metabolic models (GEMs) are knowledge-bases reconstructed from all known metabolic genes of an organism, systematically linking metabolites, reactions, and genes^26^. Integration of i-modulons with GEMs allows us to probe the interaction between the regulatory and metabolic networks. To visualize this crosstalk at the systems level, we overlaid the i-modulons onto central metabolism and amino acid metabolism pathways of the *S. aureus* metabolic reconstruction iYS854 (**Figure 3a**)^27^.

**Figure 3:**
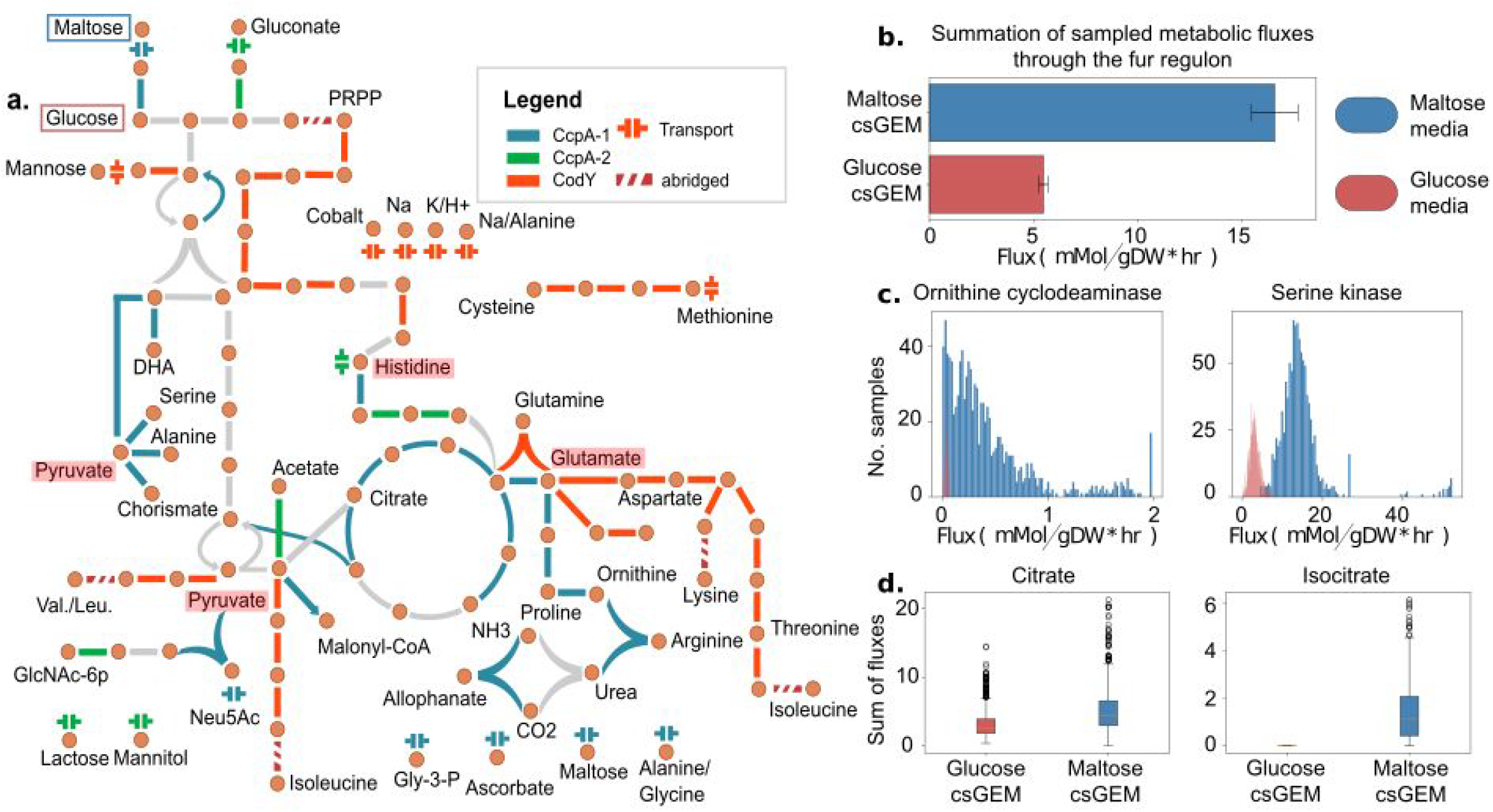
Regulation of central metabolism and its interaction with other metabolic subsystems. (a) Overlay of i-modulons onto the map of central metabolism and amino acid metabolism of *S. aureus*. The two main regulators of these metabolic subsystems, ccpA (blue/green) and codY(orange), control central carbon and nitrogen metabolism respectively. These two i-modulons intersect at key metabolic nodes-- namely pyruvate, histidine, and glutamate (highlighted in red). Entry points of sugars used in the next section, glucose and maltose, are highlighted with red and blue boxes respectively. (b) Activity of reactions associated with the Fur i-modulon in presence of different carbon sources: maltose and glucose. The proxy used here is the sum of median sampled fluxes through reactions catalyzed by enzymes which form part of the Fur i-modulon. Transcriptomic measurements indicated an unexpected increase in Fur i-modulon activity when carbon source was switched from glucose to maltose, a result which is recapitulated through metabolic modeling. (c) Reactions associated with the Fur i-modulon having the largest increase in simulated flux in glucose. (d) Calculated proxy for intracellular metabolite concentrations. Despite TCA cycle shutdown in glucose medium, the summation of fluxes through citrate is non-null while that of isocitrate is indicating that flux rewiring occurs at the citrate node.

The CcpA-1 and CodY i-modulons dominate regulation of the genes in these metabolic subsystems of *S. aureus*. The two CcpA i-modulons controlled many of the genes in carbon metabolism. The genes required for the tricarboxylic acid (TCA) cycle were found primarily in the CcpA-1 i-modulon, with the exception of genes encoding fumarase and malate dehydrogenase. Additionally, the CcpA-1 i-modulon contained genes required for degradation of gluconeogenic amino acids (serine, histidine, and alanine) and secondary metabolites (chorismate and N-acetyl-neuraminic acid). Also included were genes encoding two key gluconeogenic enzymes - phosphoenolpyruvate (PEP) carboxykinase and fructose-1,6-bisphosphatase. Genes involved in transport of alternate carbon sources were also present. In contrast to catabolic CcpA-regulated genes, the i-modulon associated with *codY* regulation was dominated by genes participating in biosynthesis of amino acids lysine, threonine, methionine, cysteine, histidine, and branched chain amino acids (BCAA) isoleucine, leucine, and valine^14^. Regulation of interconversion between glutamine and glutamate (*gltA*), a key component of nitrogen balance and assimilation, was also a part of the CodY i-modulon.

While the two i-modulons (CcpA-1, CodY) did not share any genes, they intersected at some key metabolite nodes in central metabolism, including pyruvate, glutamate, histidine, and arginine. Genes in the CcpA-1 i-modulon encode enzymes that generate pyruvate from amino acids and use the pyruvate to generate energy through fermentation, synthesize glucose via gluconeogenesis, or synthesize fatty acids via malonyl-coA. Enzymes encoded by genes in the CodY i-modulon, on the other hand, redirect pyruvate to instead synthesize BCAA. Similarly, glutamate is directed towards the urea cycle by CcpA-1 and towards biosynthesis of the aspartate family amino acids by CodY. While genes required for catabolism of histidine are in the CcpA-1 i-modulon, genes encoding histidine biosynthesis is instead part of the CodY i-modulon.

Interestingly, while CcpA regulates L-arginine synthesis from L-proline (via *putB, rocD*, and *rocF*), CodY regulated genes involved in L-arginine synthesis from L-glutamate (*argJ* and *argB*). This dichotomy, combined with the observation that isoleucine affects CodY-dependent repression^14^, explains why none of *argJBCF* were expressed in JE2 *ccpA*∷*tetL* in CDM, a BCAA-rich medium^28^. It is likely that CodY- and CcpA-mediated repression are both active in CDM. By controlling the expression of these key metabolic genes, CcpA-1 and CodY i-modulons can readily redirect the fluxes through different metabolic subsystems.

### GEMs compute a flux-balanced state that reflect regulatory actions of CcpA

Metabolic network reconstructions can be converted into genome-scale models that allow for the computation of phenotypic states^29^. We can compute the optimal flux through the metabolic network using flux-balance analysis (FBA)^27^. In particular, we can compute the metabolic state that is consistent with nutrient sources in a given environment to support optimal bacterial growth. In the previous CcpA i-modulon validation experiment, we observed that changing the carbon source from glucose to maltose in R10LB also led to an unexpected spike in activity of the iron-responsive Fur i-modulon (**Figure S3a)**. To investigate whether there was a possible metabolic role explaining the increase in Fur activity, we generated two condition-specific GEMs (csGEMs) starting with iYS854^27^. For both csGEMs, we computed the state of the metabolic network that supports growth in R10LB, with either glucose or maltose as the main carbon source (Methods and Materials). We assumed that CcpA-1 repression was active only when glucose was the main glycolytic nutrient source (and the corresponding set of reactions was shut off). Reaction fluxes across the network were then sampled using flux-balance analysis, assuming that the bacterial objective was biomass production^30^. Sampling accounts for different network flux distributions that can achieve the same optimal solutions (i.e., identical biomass production rates).

Under these conditions, the sum of sampled fluxes through reactions associated with the Fur i-modulon was significantly higher in maltose media (Kolmogorov-Smirnov test, p < 0.01, statistics > 0.9), confirming that the spike in Fur activity could be a result of metabolic flux rewiring **(Figure 3b**). In particular, fluxes through serine kinase (*sbnI*, a precursor metabolic step of staphyloferrin B biosynthesis) and ornithine cyclodeaminase (*sbnB*) were significantly increased (**Figure 3c**). These changes came as a result of flux rewiring away from deactivated metabolic steps. For example, due to arginase (*rocF*) deactivation, the flux through half of the urea cycle and ornithine cyclodeaminase was lower. Similarly, serine deaminase (*sdaB*) - located two metabolic steps downstream of serine kinase - was deactivated due to simulated down-regulation of genes in the CcpA-1 i-modulon, and flux through phosphoglycerate dehydrogenase, serine kinase, and phosphoserine phosphatase was decreased.

We computed the sum of fluxes producing each metabolite as a proxy for intracellular concentrations and found that the calculated values were significantly larger in maltose media for 68 metabolites including ammonium, glutamate, and isocitrate. The majority of the TCA cycle was shut off in the glucose-specific GEM (due to simulated repression of *citB*, *icd*, *odhA*, *sdhABCD*, and *sucCD*), and therefore the concentration proxy for isocitrate was essentially null, while that of citrate was not (**Figure 3d**). Previous studies have shown that *citB* deletion results in increased intracellular concentration of citrate ^31^. Apart from being an intermediate in the TCA cycle, citrate can be utilized in the model as a precursor to staphyloferrin A and staphyloferrin B biosynthesis (which are included in the Fur i-modulon), or it can be converted back to oxaloacetate and acetate *via* citrate lyase. All three routes were part of the solution space, with citrate lyase carrying the largest median flux. Taken together, these modeling simulations suggest that utilizing maltose instead of glucose induces metabolic flux rewiring towards reactions associated with the Fur i-modulon.

### An i-modulon details possible scope and functions of sigma factor σ^s^

Global stress response in *S. aureus* is modulated by the alternate sigma factor σ^B 32,33^. Though two other sigma factors, σ^S^ and σ^H^, have been recognized in this organism, their exact functions and full regulon are not as well understood^34,35^. We identified two i-modulons that correspond to sigma factors σ^B^ and σ^S^. The SigB i-modulon contained genes encoding σ^B^ (*sigB)*, anti-σ^B^ (*rsbW)*, and anti-σ^B^ antagonist (*rsbV)*. The activity of SigB i-modulon was correlated with *sigB* expression (Pearson R = 0.55, p-value = 8.2e-11) (**Figure S4a**), with the highest activation in stationary phase (OD_600_ = 1). Furthermore, a conserved 29 bp motif was enriched from 28 unique regulatory regions of SigB i-modulon genes (**Figure S4b**) (see Methods and Materials).

As the regulatory role of σ^B^ has been previously explored in detail^33,36–39^, we focused here on the less understood regulatory role of σ^S^. Though σ^S^ is important for both intracellular and extracellular stress response, its full regulon has yet to be defined^35,40^. ICA identified a large i-modulon with 137 genes including *sigS* itself (which encodes σ^S^). As with the SigB i-modulon, expression of the *sigS* gene correlated to activity of the ICA-derived SigS i-modulon (Pearson R = 0.77, p-value = 4.26e-22) (**Figure 4a**). Previous studies have shown that CymR represses *sigS* expression and therefore may lead to its decreased activity^41^. We confirmed this relationship as the SigS i-modulon activity was anti-correlated with the CymR i-modulon activity (Pearson R = −0.68, p-value = 8.23e-10) **(Figure S4c**).

**Figure 4:**
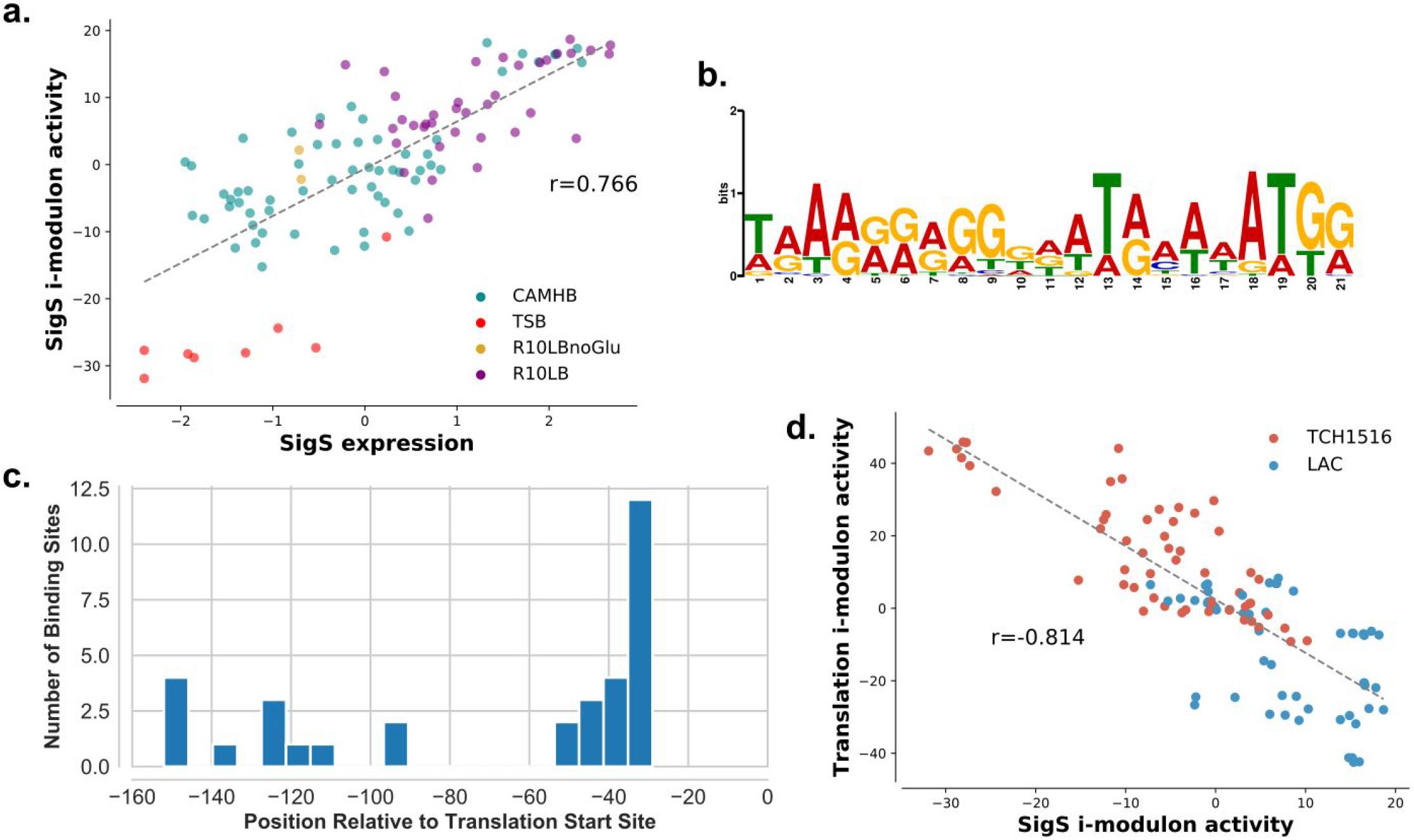
Profiling alternate sigma factor S. (a) The expression level of *sigS* gene and the activity level of the SigS i-modulon shows strong positive correlation. (b) The regulatory region (150 bp upstream of the first gene in operon) of genes in the SigS i-modulon contained a conserved purine rich motif. (c) The positions (relative to transcription start-site) of the enriched motif within the regulatory sites of genes in the SigS i-modulon. For many genes in the SigS i-modulon, the motif was present ~35 bp upstream of the translation start site. (d) ‘Greed vs. Fear’ trade-off is reflected in the activity of the Translation (greed) and SigS (fear) i-modulons. LAC showed increased propensity for fearful bet-hedging strategy while TCH1516 relied on a more greedy strategy.

To further characterize σ^S^, we looked for conserved motifs in the regulatory regions of the genes in the i-modulon and found a purine-rich 21 base-pair purine rich motif (E-value = 7.7e-8) in the regulatory region of at least 56 genes in the SigS i-modulon (**Figure 4b**). Comparisons against a known prokaryotic motif database revealed that the *S. aureus* σ^S^ motif was most similar to that of the σ^B^ (MX000071) motif in *B. subtilis* (E-value = 1.62e-02) (see Materials and Methods). Next, we analyzed the distance between the center of the motif and the transcription initiation site. For most genes, the motif was present at or around 35 base-pairs upstream of the translation start site, though motifs were also found further upstream (**Figure 4c**).

Of the 137 genes in the i-modulon, only 56 (~41%) had an assigned function in the reference genome, further highlighting our limited understanding of σ^S^ functionality. However, many of the annotated gene products were key factors in controlling cellular state. These included factors regulating virulence (*sarA, sarR, sarX*), antimicrobial resistance (*cadC, blaI*), metabolism (*arcR*, *argR*), cell wall biogenesis (*vraRST*), biofilm formation *(icaR*), and DNA damage repair (*recX)*. Genes encoding proteins critical for stress response such as universal stress protein (Usp), toxin MazF, competence proteins ComGFK, and cell division protein were also present.

The SigS i-modulon also plays a critical role in the so-called ‘fear vs. greed’ trade-off in *S. aureus*. Previously described in *E. coli*, this trade-off describes the allocation of resources towards optimal growth (greed) versus allocation towards bet-hedging strategies to mitigate the effect of stressors in the environment (fear)^7,42^. This balance is reflected in the transcriptome composition as an inverse correlation between the activities of the stress-responsive SigS i-modulon and the Translation i-modulon (Figure 4d). Unlike *E. coli*, however, this relationship was independent of growth rate, as growth rate had weak correlation with Translation i-modulon expression activity (Pearson R = 0.094, p-value = 0.514). Interestingly, mapping this trade-off highlighted a possible difference in survival strategy between the two USA300 strains. TCH1516 tended towards a greedy strategy with high Translation i-modulon activity while LAC was more likely to rely on bet-hedging, or fear.

### ICA reveals organization of virulence factor expression

ICA captured systematic expression changes of several genes encoding virulence factors. Previous studies described over half a dozen transcription factors with direct or indirect roles in regulation of virulence factor expression in *S. aureus*^*43*^. The number of regulators, and their complex network of interactions, make it extremely difficult to understand how these genes are regulated at a genome scale. In contrast, ICA identified only three i-modulons - named agr, SaeR, and Vim-3 - that were mostly composed of virulence genes (**Figure 5a**). The activity level of Agr had extremely low correlation with that of SaeR and Vim-3, suggesting that Agr may have only limited cross-talk with the other two i-modulons (**Figure S5a**). However, the activity levels of SaeR and Vim-3 were negatively correlated (Pearson R = −0.57, p-value = 8.6e-11). As the two i-modulons contain different sets of virulence factors, the negative correlation points to a shift in the virulence state where *S. aureus* may adopt different strategies to thwart the immune system. Collectively, the three virulence i-modulons revealed coordinated regulation of 65 genes across the genome. These results suggest that the complexity behind virulence regulation can be decomposed into discrete signals and the virulence state of *S. aureus* can be defined as a linear combination of these signals.

**Figure 5:**
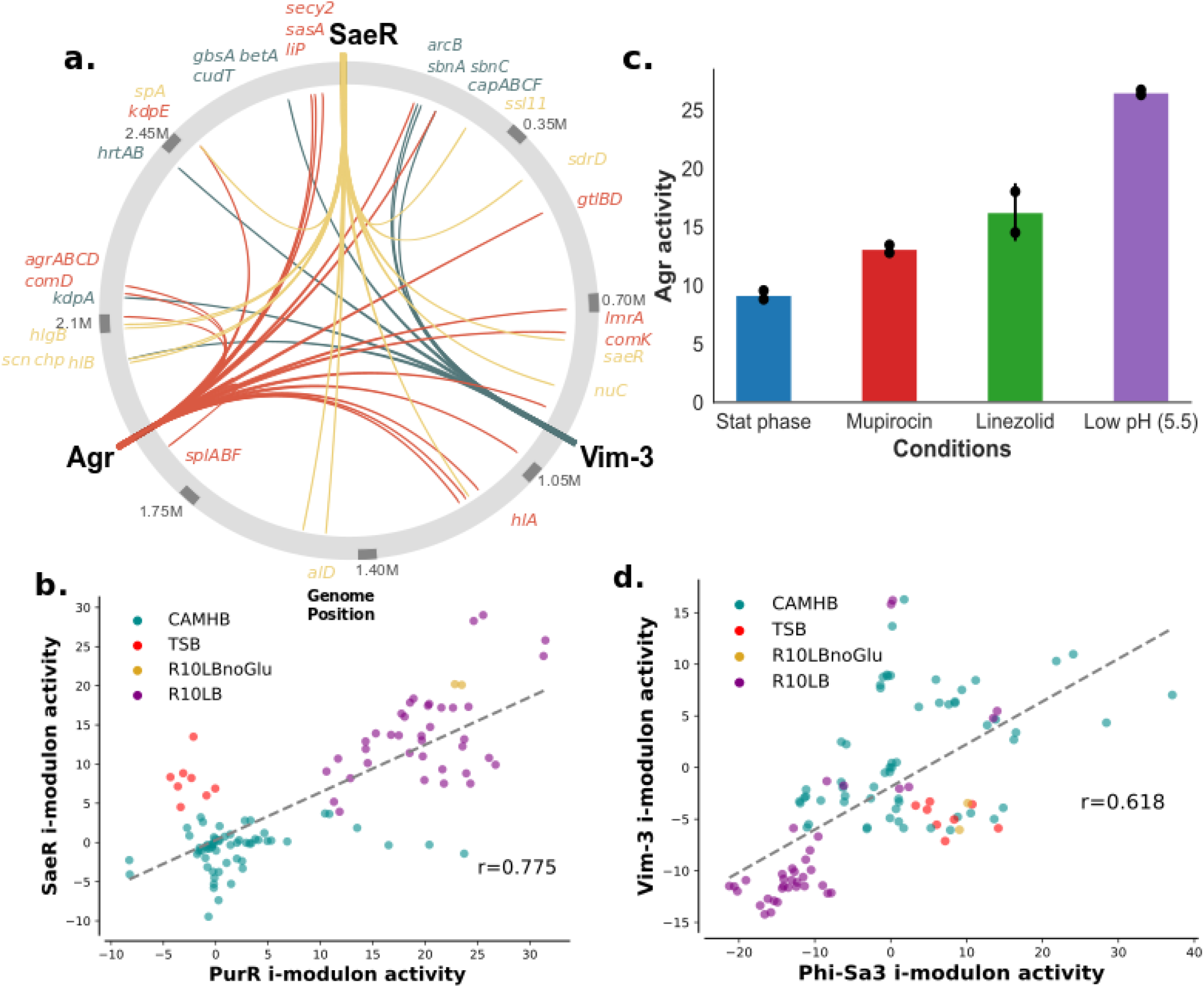
Global regulation of virulence factors. (a) The three virulence i-modulons (SaeR, Agr, Vim-3) and the genomic positions of the genes in their respective i-modulons are mapped. The signals encode over 25 virulence factor associated genes. (b) PurR i-modulon activity is highly correlated with virulence i-modulon SaeR. (c) Challenge with low pH, linezolid, and mupirocin leads to strong activation of agr in exponential growth phase. Interestingly, this activation is stronger than that induced by stationary phase (O.D. 600 = 1.0). Activation of agr was much weaker under all other experimental conditions considered. (d) Co-activation of Phi-Sa3 i-modulon with virulence i-modulon Vim-3.

The SaeR i-modulon contained 27 genes, including the genes for the SaeRS two-component system (TCS). The activity level of this i-modulon strongly correlated with the expression level of *saeRS*, further supporting the idea that the genes in this i-modulon are regulated (directly or indirectly) by SaeRS (Pearson R = 0.80, p-value = 1.38e-25). Furthermore, the virulence genes *chp, coa, ssl11, sbi, map, lukA*, and *scn*, previously reported to be under the control of SaeRS^44^, were also found in this i-modulon. PurR, a transcription factor that regulates the genes of purine biosynthesis, has been recently implicated in regulation of virulence factors^45,46^. Consistent with this observation, the activity level of the SaeR i-modulon correlated well (PearsonR = 0.77, p-value = 8.9e-23) with the activity of the PurR i-modulon (**Figure 5b**). Thus, SaeR may act as a bridge between virulence and metabolism.

Similarly, the Agr i-modulon contained the *agrABCD* genes involved in regulation of the quorum sensing agr regulon^47,48^. As most of our samples were collected during early-to mid-exponential growth phase, the Agr i-modulon remained inactive in these conditions (**Figure S5b**). Only acidic conditions (pH 5.5) and treatment with translation inhibitors linezolid and mupirocin activated agr during exponential growth (**Figure 5c**). Both pH- and translation inhibition-dependence of agr expression have been previously reported ^49–52^. Unexpectedly, the Agr i-modulon was activated to a much greater extent by these factors than high cell density (O.D. 1.0), for which its role in quorum sensing is extensively characterized.

The Vim-3 virulence i-modulon consisted of genes required for siderophore and heme utilization (*sbnABC, hrtAB)*, capsule biosynthesis (*cap8a, capBC, cap5F*), and osmotic tolerance (*kdpA, betAT, gbsA*). The Vim-3 i-modulon had maximal activity under hyperosmotic condition introduced by 4% NaCl and when grown to stationary phase (OD 1.0) in CAMHB (**Figure S5c**). The increased expression of capsule biosynthesis genes have shown to be responsive to change in osmotic pressure as well as iron starvation, which is consistent with the inclusion of iron scavenging and osmotic tolerance genes in the i-modulon with the capsular biosynthesis genes^53,54^.

We further identified a prophage Phi-Sa3 associated i-modulon as a new putative i-modulon required for virulence. The Phi-Sa3 i-modulon consists of genes in the Phi-Sa3 prophage and several genes encoding DNA replication and repair enzymes. Excluded from the i-modulon were the virulence factors that were horizontally acquired along with the phage (*scn and chp*)^55^, which now fell under the control of SaeR. Of the four phages in *S. aureus* strain Newman, Phi-Sa3 is the only prophage that is unable to generate complete viral particles when challenged with DNA damaging agent mitomycin^56^. However, evidence suggests that this prophage is still active in USA300 strains and its genes are expressed during lung infection, where it may play a role in establishing virulence^57^. Corroborating this hypothesis, we found that the activity of the Phi-Sa3 i-modulon correlated highly with the Vim-3 i-modulon (Pearson R = 0.62, p-value = 9.9e-13)(Fig. 5e). As the Phi-Sa3 i-modulon does not contain any virulence genes, the phage itself may play an accessory role in establishing virulence.

### ICA model provides a platform for ex-vivo data interpretation

Transcriptomic models based on ICA can also be used to interpret new expression profiling data. Any new data, even those from *ex-vivo* conditions, can be projected onto the i-modulon structure of the TRN to convert the values from gene expression levels to i-modulon activity levels (see Materials and Methods). Here we show that the projection of *ex-vivo* data on to the model recapitulates expected results and provides new insights into *S. aureus* transcriptional changes in human blood.

We projected previously published time course microarray data collected from *S. aureus* USA300 LAC grown in Tryptic Soy Broth (TSB), human blood, and serum^58^. Bacteria grown to exponential phase in TSB was used as inoculum for all samples; we used this as our new base condition for the projected data. Therefore, all i-modulon activity levels in this set represents log2 fold change in activity from this base condition. Once transferred to serum, the activities of Fur and CodY i-modulons in serum increased dramatically (**Figure 6a**). The large change in activity coupled with the sizeable number of genes in each i-modulon (80 and 45 genes in CodY and Fur, respectively) indicates that *S. aureus* reallocates a considerable portion of its transcriptome to reprogram amino acid and iron metabolism in serum. PurR and SaeR activity also increased, though their magnitude of change was dwarfed by the changes in activity of CodY and Fur. Agr activity, on the other hand, declined and remained low over the two hour period. Because *agr* positively regulates a number of virulence genes, its activity level may be expected to go up in serum. However, consistent with the model prediction, *agr* regulated genes have been shown to be transcriptionally down-regulated in serum due to sequestration of the *agr* activator, auto-inducing peptides (AIP), by human serum component apolipoprotein B ^59,60^.

**Figure 6:**
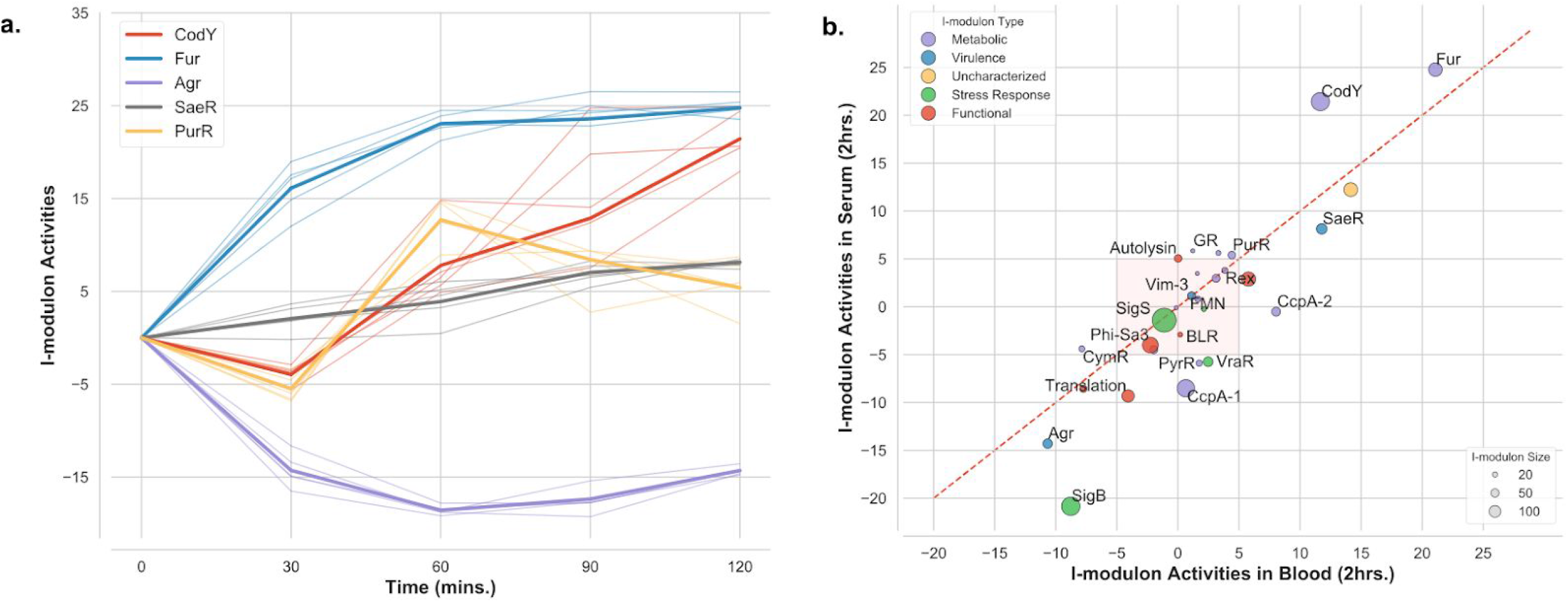
ICA analysis of blood and serum data. (a) Activity levels of select i-modulons in serum over the two hour time period. The thick line represents the mean activity across all replicates and the thin line represents activity in each individual replicate (n=4). Activity levels were around the inoculum values, Comparison of i-modulon activity between serum and blood and two hour time point. The dashed red line is the 45 degree line; i-modulons below the line have higher activity in blood and those above the line have higher activity in serum. Red shaded area contains i-modulons with less than 5 fold change in activity in both conditions.

We next calculated the differences in i-modulon activities in blood and serum at the final two hour time-point. Fur, CodY, PurR, SaeR, and Agr had similar activity levels in both blood and serum (**Figure 6b**). Therefore, the activity of these regulators are likely governed by the non-cellular fraction of the blood. I-modulons PyrR, SigB, Translation, VraR, CcpA-1, and CcpA-2 had higher activity levels in blood than in the serum (**Figure 6b**). Glucose concentration in blood is lower than in serum, which likely explains the shift in CcpA-1 activity^61^. The lower glucose concentrations relieves CcpA mediated repression of its regulon, leading to higher expression. The shift in the PyrR i-modulon also corroborates previous studies, which demonstrated that *S. aureus* strain JE2 (a derivative of LAC) requires more pyrimidine when growing in blood than in serum^62^. The signals or cues driving the change in activity of the other i-modulons (SigB, Translation, VraR, and CcpA-2) remains unknown. Overall, our analysis revealed that during acute infection, CodY and Fur play key roles in rewiring the *S. aureus* metabolism in serum and blood when compared to TS,B while SaeR (and not Agr) drives the virulence gene expression. In addition, SigB, Translation, and VraR i-modulons are uniquely activated by the cellular fraction of the blood and may thus be responding to unique stresses they impart.

## Discussion

Here, we described an ICA-based method to elucidate the organization of the modules in TRN in *S. aureus* USA300 strains. Using this method, we identified 29 independently modulated sets of genes (‘i-modulons’) and their activities across the sampled conditions. This framework for exploring the TRN provides three key advantages over traditional methods, especially when working with non-model organisms: (1) the method provides an quantitative reconstruction of the TRN; (2) it is an untargeted, and therefore unbiased, approach; and (3) the approach utilizes expression profiling data, an increasingly ubiquitous resource.

First, i-modulons quantitatively capture the complexities of transcriptional regulation and enable a new way to systematically query the transcriptome. By recasting the data in terms of explanatory i-modulons, we gained a deeper understanding of large changes in transcription profiles between CAMHB bacteriologic media and the more physiologically relevant mammalian tissue culture-based media R10LB. The analysis reduced the number of features needed to capture most of the information in the transcriptome from hundreds of genes to fifteen i-modulons. Additionally, quantified activity levels of i-modulons also enabled integration of regulatory activity with metabolic models and revealed coordination between metabolic and regulatory networks. Such reduction in complexity and the integration of different aspects of *S. aureus* biology (e.g., virulence, metabolism, stress response, etc.) will be crucial to understanding the mechanisms that enable successful infection *in-vivo*.

Second, this method presents a platform for untargeted, global analysis of the TRN. Due to its untargeted nature, we also identified two key virulence features of *S. aureus*. ICA revealed coordinated regulation of genes in capsule biosynthesis, osmotic tolerance, and iron starvation (Vim-3 i-modulon). Both capsule formation and siderophore scavenging are important in nasal colonization^63,64^. Similarly, growth of *S. aureus* in Synthetic Nasal Medium (SNM3) increases the expression of genes required for osmotolerance. Therefore, the Vim-3 i-modulon may represent a concerted regulation of genes required for successful nasal colonization. We also identified the Phi-Sa3 phage i-modulon, whose activity level correlated with that of the Vim-3 i-modulon. The Phi-Sa3 i-modulon did not include the virulence genes (e.g. *sak, scn*, etc.) that were acquired with the phage, suggesting that phage replication genes were expressed independently of the virulence genes. Given that its activity was correlated with Vim-3, this phage may also play an important role in nasal colonization.

Lastly, ICA uses RNA-seq data to extract information about the TRN, making it more accessible to non-model organisms including *S. aureus*. Reconstructing the TRN with traditional methods is highly resource intensive, as they require targeted antibodies or specialized libraries of plasmids containing all transcription factors of interest^65^. While these approaches have given us great insights into TRNs of model organisms like *E. coli^10^*, such comprehensive data is not available for most microbes. Several studies have attempted to circumvent this by comparing the expression profiles of wild-type *S. aureus* strains with their counterpart through transcription factor knockout or introduction of a constitutively active transcription factor^66,67^. However, these approaches often overestimate the regulatory reach of the transcription factor, as such genetic changes can trigger the differential expression of genes not directly under the regulator’s control. In contrast, the ICA-based method uses increasingly ubiquitous RNA-seq data combined with machine learning to identify and characterize i-modulons. While a large amount of data is required to build such a model, an abundant number of expression profiles (e.g., 3530 samples from multiple *S. aureus* strains) are already publicly available on Gene Expression Omnibus (GEO). Indeed, utilizing only RNA-seq data, we predicted the previously unknown regulon of stress-associated sigma factor σ^S^ and its possible roles in biofilm formation, growth rate control, and general stress response. With the growing number of available expression profiles, such characterizations can be extended to other undefined or poorly defined regulons. Therefore, in the absence of a comprehensive set of targeted antibodies against *S. aureus* transcription factors, reanalyzing the publicly available database with ICA could be used to further reconstruct its TRN.

We have shown that ICA based decomposition can be utilized to build a quantitative and explanatory model of *S. aureus* TRN from RNA sequencing data. Application of this model enabled us to query metabolic and regulatory crosstalk, discover new potential regulons, find coordination between metabolism and virulence, and unravel the *S. aureus* response to *ex-vivo* conditions. Due to this versatility this model and other models generated through this framework will be a powerful tool in any future studies of *S. aureus*.

## Materials and Methods

### RNA extraction and library preparation

*S. aureus* USA300 isolates LAC, TCH1516 and ALE derivatives of TCH1516 (SNFR and SNFM) were used for this study. The growth conditions and RNA preparation methods for data acquired from Choe et al. has been previously described^68^. Detailed growth conditions, RNA extraction and library preparation methods for other samples have also been already described^69^. Briefly, an overnight culture of *S. aureus* was used to inoculate a pre culture and were grown to mid-exponential growth phase (OD600 = ~0.4) in respective media (CAMHB, RPMI + 10% LB, or TSB). Once in mid-exponential phase, the preculture was used to inoculate the media containing appropriate supplementation or perturbations. Samples were collected at O.Ds and time-points indicated in the metadata (Supplementary File 1). All samples were collected in biological duplicates originating from different overnight cultures. Sample for control conditions were collected for each set to account for batch effect.

### Determining Core Genome with Bi-Directional BLAST Hits (BBH)

To combine the data from the two strains, core genome containing conserved genes between the LAC (GenBank: CP035369.1 and CP035370.1) and TCH1516 (GenBank: NC_010079.1, NC_012417.1, and NC_010063.1) were first established using BBH^70^. In this analysis, all protein sequences of CDS from both genomes are BLASTed against each other twice with each genome acting as reference once. Two genes were considered conserved (and therefore part of the core genome) if (1) the two genes have the highest alignment percent to each other than to any other genes in the genome, (2) the alignment percent is at least 80% and (3) the coverage is at least 80%.

### RNA Sequencing Data Processing

The RNA sequencing pipeline used to analyze and perform QC/QA has been described in detail previously^69^. Briefly, the sequences were aligned to respective genomes, LAC or TCH1516 (NC_010079.1, NC_012417.1, NC_010063.1) using Bowtie2^71,72^. The samples from ALE derivatives, SNFM and SNFR, were aligned to TCH1516. The aligned sequences were assigned to open reading frames using HTSeq-counts ^73^. Differential expression analysis was performed using DESeq2 with p-value threshold of 0.05 and an absolute fold change threshold of 2^74^.

To create the final counts matrix, counts from conserved genes in LAC samples were represented by the corresponding ortholog in TCH1516. The counts for accessory genes were filled with 0s if the genes were not present in the strain (i.e. LAC specific genes had counts of 0 in TCH1516 samples and vice versa). Finally, to reduce the effect of noise, genes with average counts per sample < 10 were removed. The final counts matrix with 2581 genes was used to calculate Transcripts Per Million (TPM).

### Computing robust components with ICA

Procedure for computing robust components with ICA has been described in detail previously^7^. Log_2_(TPM + 1) values were centered to strain specific reference conditions and used as input of ICA decomposition. These conditions are labeled: ‘USA300_TCH1516_CAMHB_U01-Set000_Control_1’, ‘USA300_TCH1516_CAMHB_U01-Set000_Control_2’ for TCH1516 and ‘USA300_LAC_CAMHB_U01-Set001_Control_1’, ‘USA300_LAC_CAMHB_U01-Set001_Control_2’ for LAC.

Next, Scikit-learn (v0.19.0) implementation of FastICA algorithm was used to calculate independent components with 100 iterations, convergence tolerance of 10^−7^, log(cosh(x)) as contrast function, and parallel search algorithm^75,76^. The number of calculated components were set to the number of components that reconstruct 99% of variance as calculated by principal component analysis.

The resulting S-matrices containing source components from the 100 iterations were clustered with Scikit-learn implementation of DBSCAN algorithm with epsilon of 0.1, and minimum cluster seed size of 50 samples (50% of the number of random restarts). If necessary, the component in each cluster was inverted such that the gene with the maximum absolute weighting the component was positive. Centroids for each cluster was used to define the final weightings for **S** and corresponding **A** matrix.

The whole process was repeated 40 times to ensure that the final calculated components were robust. Finally, components with activity levels that deviated more than 5 times between samples in the same conditions were also filtered out.

### Determining independently modulated sets of genes

ICA enriches components that maximize the non-gaussianity of the data distribution. While most genes have weightings near 0 and fall under gaussian distribution in each component, there exists a set of genes whose weightings in that component deviates from this significantly. To enrich these genes, we used Scikit-learn’s implementation of the D’Agostino K^2^ test, which measures the skew and kurtosis of the sample distribution^77^. We first sort the genes by the absolute value of their weightings and perform the K^2^ test after removing the gene with the highest weighting. This was done iteratively, removing one gene at a time, until the K^2^ statistic falls below a cutoff.

We calculated this cutoff based on sensitivity analysis on agreement between enriched i-modulon genes and regulons inferred by RegPrecise^78^. For a range of cutoff (between 200-600), we ran the iterative D’Agostino K^2^ test on all components and checked for statistically significant overlap of i-modulons with the regulons predicted by RegPrecise using Fisher’s Exact Test. For i-modulons with significant overlap, we also calculated precision and recall. The cutoff of 280 which led to the highest harmonic average between precision and recall (F1-score) was chosen as the final cutoff.

### Designating biological annotations to i-modulons

To designate proper annotations to i-modulons, we first compiled a dataset containing previously predicted features such as regulons, genomic islands and plasmids. The regulons in the datasets were inferred by either RegPrecise algorithm and by RNA-seq analysis of transcription factor knockout strains or strains with constitutively active transcription factors(see **Supplementary Table 2**)^66,68,79–81^. Genomic islands were determined by online IslandViewer4 tool^82^ and phages were identified with PHASTER^83^. For studies using different strains of *S. aureus* orthologs for TCH1516 and LAC were determined using BBH. The enriched genes in i-modulons were compared against this dataset for significant overlap using Fisher’s Exact Test with FDR of 10^−5^. With this analysis 15 i-modulons were enriched with high confidence (precision >= 0.5, recall >= 0.2) and 7 were enriched with low confidence. Additionally, i-modulons containing genes with shared functions (e.g. Translation and B-lactam Resistance) were annotated manually (see Supplementary Table 3).

### Differential Activation Analysis

Distribution of differences in i-modulon activities between biological replicates were first calculated and a log-norm distribution was fit to the differences. In order to test statistical significance, absolute value of difference in activity level of each i-modulon between the two samples were calculated. This difference in activity was compared to the log-normal distribution from above to get a p-value. Because differences and p-value for all i-modulons were calculated, the p-value was further adjusted with Benjamini-Hochberg correction to account for multiple hypothesis testing problem. Only i-modulons with change in activity levels greater than 5 were considered significant.

### Motif Enrichment and Comparison

Genes were first assigned to operons based on operonDB^84,85^. For i-modulon specific motif enrichments, 150 base pairs segment upstream of all the genes in the i-modulons were collected. To avoid enriching ribosome binding sites, the segment started from 15 base pairs upstream of the translation start site. For genes in minus strand, the reverse complement of the sequence was used instead. If genes were part of an operon, then only the segment in front of the first gene in the operon was used. Motifs and their positions were enriched from these segments using the online Multiple Em for Motif Elicitation (MEME) algorithm^86,87^. The following default parameters were used: -dna -oc -mod zoops -nmotifs 3 -minw 6 -maxw 50 -objfun classic -revcomp -markov_order 0. Enriched motifs were compared to combined prokaryotic databases- CollecTF, Prodoric (release 8.9), and RegTransBase (v4) using TomTom^88–91^. The parameters for TomTom were as follows: -oc -min-overlap 5 -mi 1 -dist pearson -evalue -thresh 10.0.

### Metabolic modeling

We modeled growth in RPMI supplemented with iron, manganese, zinc, and molybdate by setting the lower bound to the corresponding nutrient exchanges in iYS854 to −1 mmol/gDW/hr (the negative sign is a modeling convention to allow for the influx of nutrients) ^27^, and −13 mmol/gDW/hr for oxygen exchange (as measured experimentally). Additionally, to account for the utilization of heme by *S. aureus* terminal oxidases, we removed heme A from the biomass reaction and added as a reactant in the cytochrome oxidase reaction with the stoichiometric coefficient obtained from the biomass reaction^92^. Next, we constructed two condition-specific GEMs (csGEMs) to compare two conditions with: 1) D-glucose as the main glycolytic source and; 2) maltose as an alternative carbon source. In the first condition, we set the lower bound to D-glucose exchange to −50 mmol/gDW/hr. Assuming that in the presence of D-glucose, *ccpA* mediates the repression of multiple genes ^23,28^, we set the upper and lower bounds of the reactions encoded by genes of the *ccpA* i-modulon to 0. Specifically, we only turned off the set of 44 reactions obtained by running the “cobra.manipulation.find_gene_knockout_reactions()” command from the cobrapy package ^93^, feeding it the model and the 52 modeled genes which form part of the *ccpA* I-modulon. As such, we implemented a method similar to the switch-based approach ^94,95^, in which the boolean encoding for the gene-reaction-rule is taken into account (*i.e.* isozymes, and protein complexes). Shutting down all of the reactions yielded a model which could not simulate growth. We thus gap-filled the first csGEM with one reaction (AcCoa carboxylase, involved in straight chain fatty acid biosynthesis). To simulate the second condition in which maltose serves as the main glycolytic source, we set the lower bound of maltose exchange to −50 mmol/gDW/hr and blocked D-glucose uptake. No regulatory constraints were added. Flux-balance analysis was implemented with the biomass formation set as the functional network objective, and fluxes were sampled in both csGEMs 1000 times using the “cobra.sampling.sample” command. To normalize flux values across conditions, we divided all fluxes by the simulated growth rate. We compared the flux distribution of each reaction in the two csGEMs using the Kolmogorov-Smirnov nonparametric test, yielding 93 reactions with significantly differing flux distributions (p-value < 0.001) having a statistic larger than 0.99. To identify whether there is a metabolic basis for the difference the Fur i-modulon stimulation between conditions, we identified a set of 34 reactions encoded by the 41 modeled genes which are part of the Fur i-modulon (again using the switch-based approach).

### Targeted High-Performance Liquid Chromatography (HPLC)

For glucose detection, samples were collected every 30 minutes and filtered as described above. Growth media was syringe-filtered through 0.22 μm disc filters (Millex-GV, MilliporeSigma) to remove cells. The filtered samples were loaded onto a 1260 Infinity series (Agilent Technologies) high-performance liquid chromatography (HPLC) system with an Aminex HPX-87H column (Bio-Rad Laboratories) and a refractive index detector. The system was operated using ChemStation software. The HPLC was run with a single mobile phase composed of HPLC grade water buffered with 5 mM sulfuric acid (H2SO4). The flow rate was held at 0.5 mL/minute, the sample injection volume was 10 uL, and the column temperature was maintained at 45°C. The identities of compounds were determined by comparing retention time to standard curves of glucose. The peak area integration and resulting chromatograms were generated within ChemStation and compared to that of the standard curves in order to determine the concentration of each compound in the samples.

### Microarray data analysis and projection

All microarray data was downloaded from GEO repository GSE25454 and processed with the Affy package in R to get gene expression level^58,96^. This dataset consists of microarray data from samples grown to exponential phase in TSB (TSB 0hr) and transferred to either blood, serum or TSB. Samples were then collected every 30 mins for 2 hours. The data was centered on ‘TSB 0 hr’ time-point. Data projection was used to convert gene expression values to i-modulon activity level as described before^7^.

## Data and Code Availability

All RNA-seq data have been deposited to the Short Read Archive (SRA). Any RNA-seq data not currently available publicly will be added before publication. The data accession numbers can be found in Supplementary Table 1. Both glucose and maltose models and their sampled fluxes are also available as supplementary material (glucose_model.json, maltose_model.json, sample_glucose.csv, sample_maltose.csv). Custom code of ICA analysis can be found on github (https://github.com/SBRG/precise-db).

## Acknowledgements

This research was supported by NIH NIAID grant (1-U01-AI124316).

## Author Contributions

SP performed RNA-seq and ICA analysis and wrote the manuscript.

HT collected samples for RNA-seq and validation experiments.

YS performed analysis with metabolic models and wrote the manuscript.

AS performed ICA analysis and contributed to the manuscript.

RS and SX performed all RNA extraction and sequencing.

HM collected samples for ALE strains SNFR and SNFM.

CO collected all the HPLC samples.

AA measured respiration rates used to constrain metabolic models.

JP, VN and BOP designed the study and contributed to the manuscript.

