## Supplementary Figures for "Revealing 29 sets of independently modulated genes in *Staphylococcus aureus*, their regulators and role in key physiological responses"

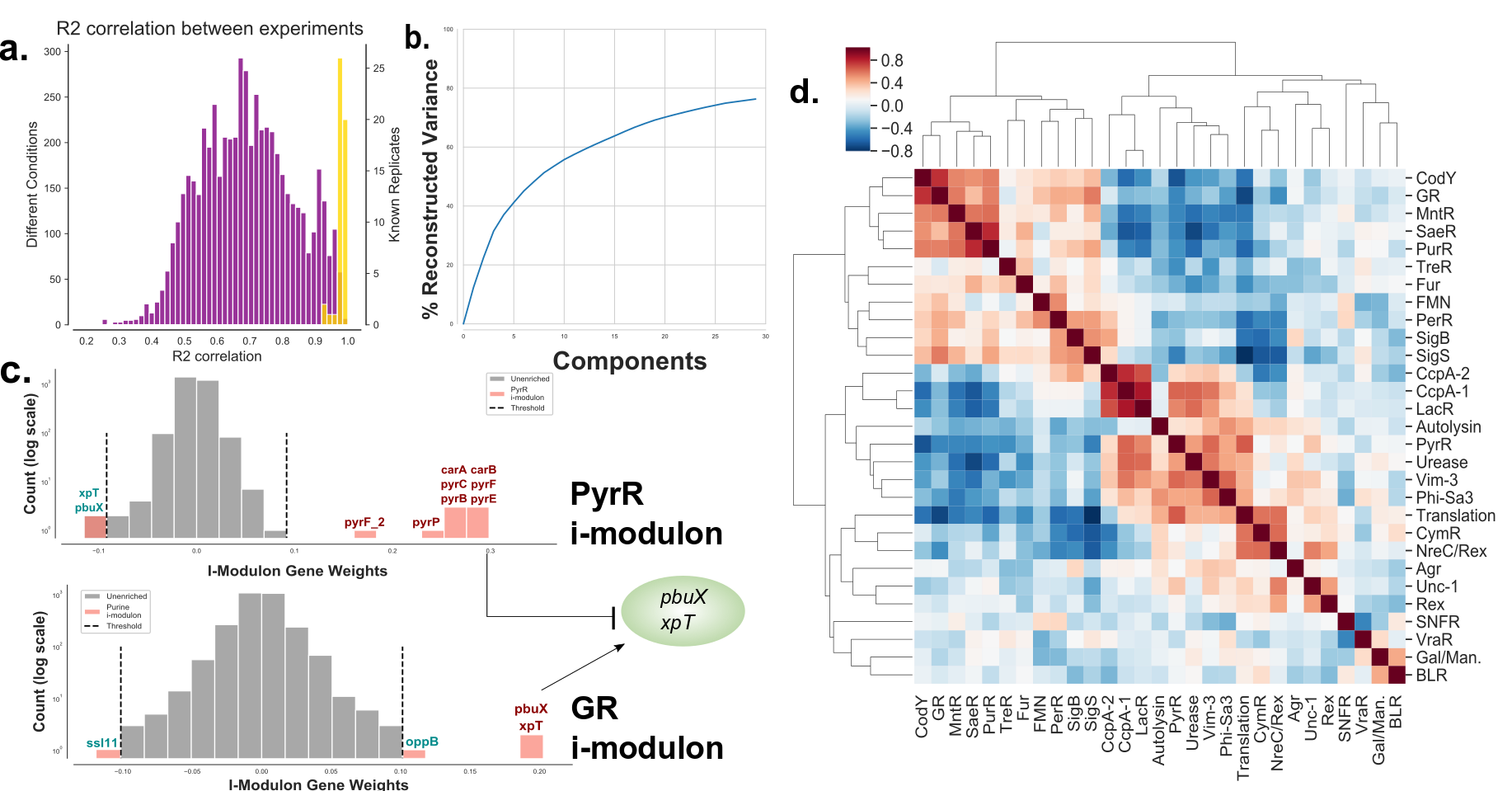

**Supplementary Figure 1: Mathematical representation of *S. aureus* TRN.** (a) RNA sequencing data were collected in duplicates and their reproducibility was verified using Spearman correlation of TPM values . Correlation between replicates (yellow bars) for all samples had R^2^ > 0.9, with most samples having R^2^ > 0.95. Correlation between different samples (purple bars) had a wide range of correlation, indicating the presence of diverse expression states. (b) The ICA decomposition captured most of the information in the input RNA sequencing compendium (**Supplementary Table 5**). 76% of the total variance could be reconstructed from the product of **S** (**Supplementary Table 6)** and **A** (**Supplementary Table 7**). (c) Histogram of gene coefficient in two example components (containing i-modulon for pyrmindine above and GR below). While most genes in a component have weights close to 0, few statistically significant outliers (outside of the vertical dashed lines) with high weightings (red bars) form an independently modulated set of genes (called an i-modulon). Genes can have both positive and negative coefficients and can be present in multiple i-modulons. The genes *xpt* and *pbuX* have negative coefficient in the PyrR i-modulon (top histogram) indicating that these genes are contra-regulated to genes with positive coefficient in the same i-modulon (.e.g *carAB*). *Xpt* and *pbuX* are also present in the GR i-modulon (bottom histogram), indicating that these two genes are regulated by multiple regulators. The first row of the matrix also contains the threshold used to call i-modulons. (d) Though i-modulons represent independently regulated set of genes, their activities are coordinated with one another. The coordination is visualized as a heatmap depicting Pearson correlation of i-modulon activities across all 108 samples.

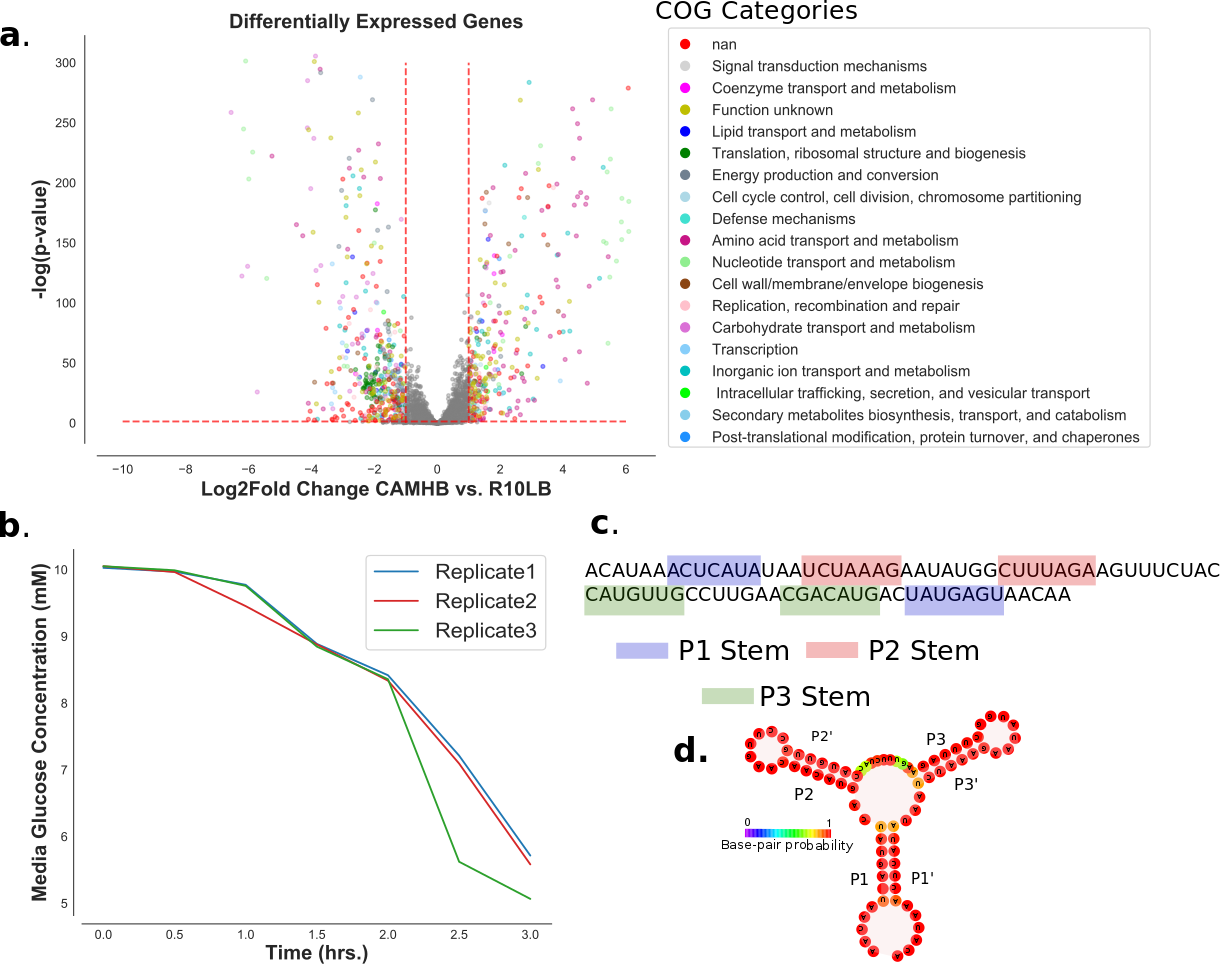

**Supplementary Figure 2: Differential activation analysis and verification.** (a) Volcano plot of differential expression levels of genes between CAMHB and R10LB. 848 genes spanning at least 17 COG categories (as determined by EggNog v4.5) were significantly differentially expressed^1^. Genes with greater than 2-fold change in expression and with p-value < 0.05 were considered significantly differentially expressed. (b) Glucose uptake was measured in R10LB and CAMHB. *S. aureus* actively took up glucose in R10LB while no glucose was detected in CAMHB. Each line represents a biological replicate in R10LB. (c) Riboswitch in conserved sequence upstream of *xpt* gene was verified using RiboSwitch^2^. (d) The structure of the riboswitch was verified with RNAfold within the ViennaRNA Package 2.0^3^.

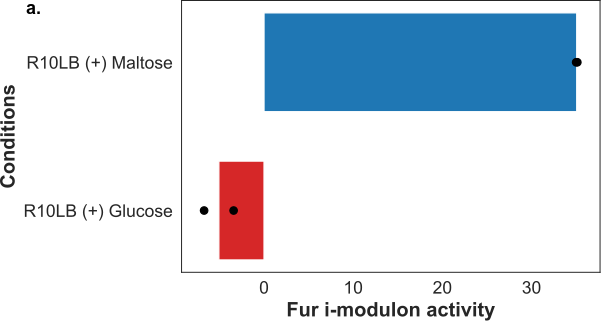

**Supplementary Figure 3: Fur activity in response to changes in carbon source.** (a) The activity of Fur i-modulons increased when the carbon source in R10LB was changed from glucose to maltose.

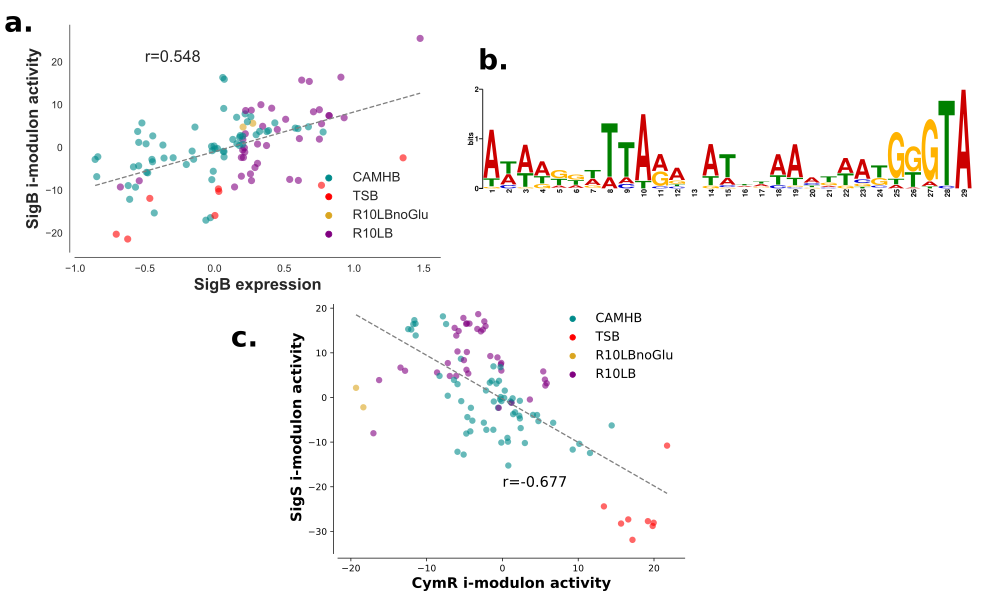

**Supplementary Figure 4: Sigma Factor I-modulons.** (a) The activity level of SigB i-modulon correlated with expression level of *sigB* gene (PearsonR=0.548, p-val=8.28e-16). (b) Regulatory region of SigB i-modulon contained a conserved motif that closely matched SigB motif of B. *subtilis*. (c) The activity level of SigS i-modulon was negatively correlated with the activity of CymR i-modulon (PearsonR=0.677, p-val=8.29e-16).

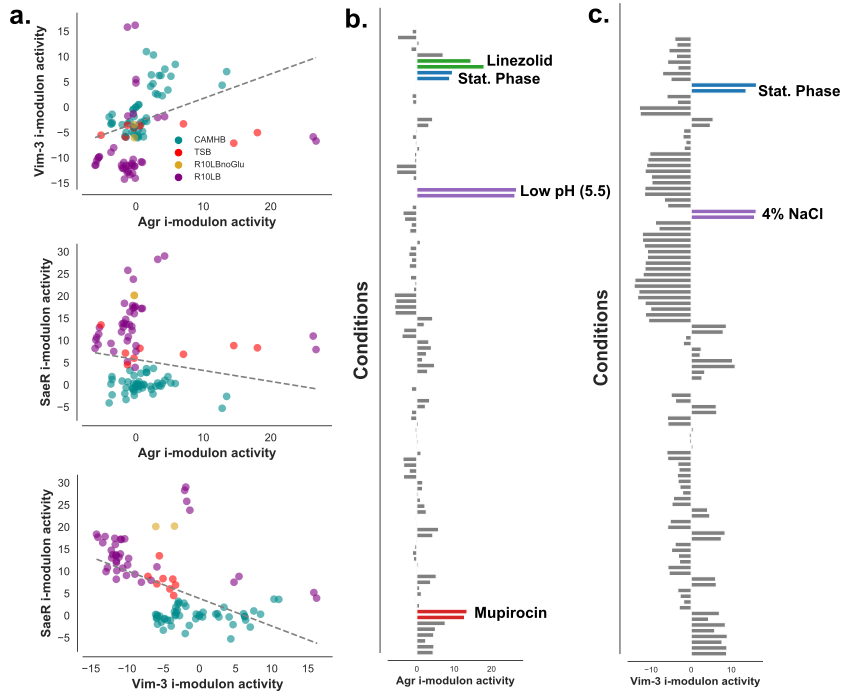

**Supplementary Figure 5: The virulence i-modulons of *S. aureus*.** (a) Activity of Agr was poorly correlated with the activities of the other two virulence associated i-modulons, SaeR and Agr. However, SaeR and Vim-3 activities were negatively correlated. (b) Agr activity in most samples were close to 0. Its activity could be induced by translation inhibitors, growth to OD600 of 1 (Stat. Phase), and low pH (5.5). (c) The Vim-3 i-modulon had the highest activity in stationary phase and when *S. aureus* was challenged with 4% NaCl.
